# Patterned apoptosis has an instructive role for local growth and tissue shape regulation in a fast-growing epithelium

**DOI:** 10.1101/2022.03.11.484029

**Authors:** Alexis Matamoro-Vidal, Tom Cumming, Anđela Davidović, Romain Levayer

## Abstract

What regulates organ size and shape remains one of the fundamental mysteries of modern biology. So far, research in this area has primarily focused on deciphering the regulation in time and space of growth and cell division, while the contribution of cell death has been much more neglected. This includes studies of the *Drosophila* wing imaginal disc, the prospective fly wing which undergoes massive growth during larval stage, and represents one of the best characterised systems for the study of growth and patterning. So far, it has been assumed that cell death was relatively neglectable in this tissue and as a result the pattern of growth was usually attributed to the distribution of cell division. Here, using systematic mapping and registration combined with quantitative assessment of clone size and disappearance, we show for the first time that cell death is not neglectable, and outline a persistent pattern of cell death and clone elimination in the disc. Local variation of cell death is associated with local variation of clone size, pointing to an impact of cell death on local growth which is not fully compensated by proliferation. Using morphometric analyses of adult wing shape and genetic perturbations, we provide evidence that patterned death affects locally and globally adult wing shape and size. This study describes a roadmap for accurate assessment of the contribution of cell death to tissue shape, and outlines for the first time an important instructive role of cell death in modulating quantitatively local growth and the morphogenesis of a fast-growing tissue.

## Introduction

The search of cellular mechanisms underlying variation in organ’s size and shape is essential to our understanding of health and evolution. These mechanisms include changes in cell shape, cell proliferation, oriented cell division, oriented cell intercalation and also cell death [1]. Programmed cell death and apoptosis are indeed essential regulators of development and morphogenesis [2]. For instance, cell death is required to eliminate scaffolding tissues which are not present in the adult body. Apoptosis in epithelia is also a driving force of morphogenesis which can help to fuse tissues by generating pulling forces [3] or trigger fold formations by generating apico-basal traction forces [4, 5]. Finally, apoptosis has been associated with the buffering of developmental fluctuations which can eliminate miss-specified/miss-patterned cells through cell competition [6], or more recently, with the buffering of mismatch between the size of the tissue and the shape of morphogen gradients [7]. Interestingly, while the functions of apoptosis are numerous, its contribution to the regulation of organ size and growth, especially in tissues undergoing fast expansion, has been poorly studied. Accordingly, emphasis has mostly been given to cellular growth regulation including the increase of cell volume and cell proliferation [8–10]. Moreover, systematic and quantitative evaluations of the pattern and number of cell deaths remain relatively rare and tedious, especially in tissues which are not appropriate for long term live imaging [11]. As such, the exact pattern of cell death and its real contribution to the regulation of organ size and shape remain poorly documented in many situations. This also applies to the *Drosophila* wing imaginal disc, one of the best studied systems for understanding patterning and growth regulation [10, 12].

*Drosophila* wing imaginal discs are epithelial sacs composed of two epithelia: the peripodial cells (squamous epithelium) and a pseudostratified epithelium that will form the adult structures [13]. The disc is separated into domains that will give rise to the adult wing (the wing pouch, or wing proper) and a distal region called the hinge which will connect the wing to the thorax. This wing pouch is patterned in different domains prefiguring adult wing structures: the pro-vein domains L2 to L5 and the dorsal-ventral boundary that will give rise respectively to the adult wing veins and the adult wing boundaries (**Figure 1A,B**). Wing imaginal discs undergo massive growth during larval development through several rounds of cell division (9 to 11 cycles) [14]. This growth phase is followed by profound remodeling and morphogenesis including tissue folding, elongation, and apposition of the two epithelial layers leading to the formation of the final adult wing shape (**Figure 1A,B**) [15], all this performed with a high degree of precision and reproducibility [16]. Decades of work sought to dissect the mechanisms controlling growth and size of the *Drosophila* wing, and have mostly focused on the distribution and regulation of cell proliferation while neglecting cell death [10]. Indeed, seminal works more than 20 years ago found that cell death is relatively minor in the wing disc and that it occurs sporadically without any noticeable pattern [17]. Yet, this did not exclude more subtle functions of cell death.

**Figure 1:**
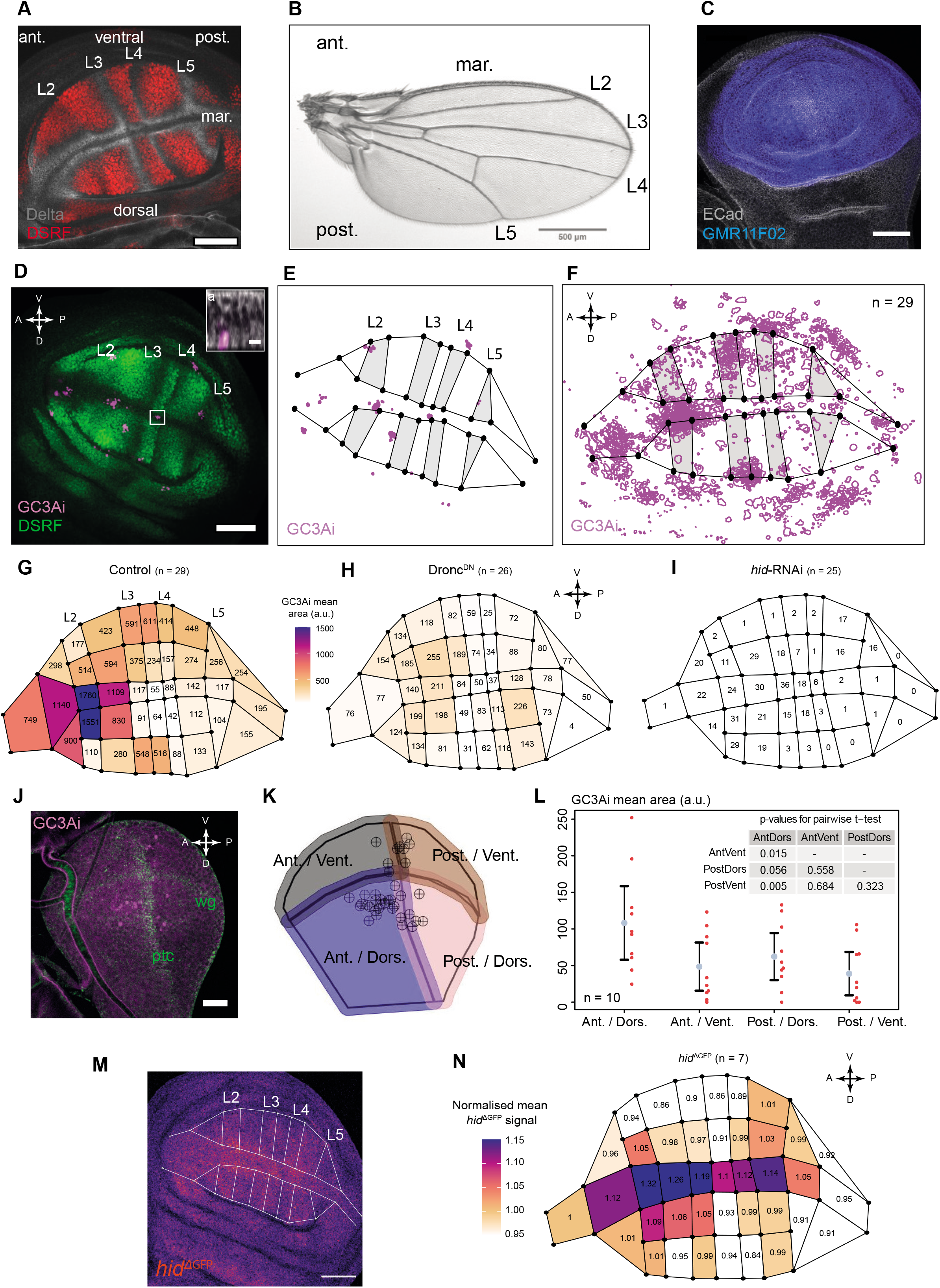
caspase-3 activity is spatially biased in the developing wing tissue. **A:** Wing disc at ~ 96 h after egg laying (AEL) showing veins (L2 to L5), interveins (red), dorsal, ventral, margin (mar.), anterior (ant.) and posterior (post.) territories. Red: anti-DSRF immunostaining. Grey: anti-Delta immunostaining. Orientation is the same for all wing discs images and schemes of this article, with the anterior compartment leftwards, and the dorsal compartment downwards. **B:** Female adult wing on dorsal view, with longitudinal veins (L2 to L5), margin (mar.), anterior (ant.) and posterior (post.) territories. **C:** Wing disc at ~ 96 h AEL showing the GMR11F02-GAL4 expression domain (blue). Gray: anti E-Cadherin immunostaining. **D:** z-projection of a wing disc showing caspase-3 activity using the GC3Ai reporter (magenta) expressed under the control of GMR11F02-GAL4 at ~ 96 h AEL. The inset is an orthogonal view of the white boxed area, showing that the GC3Ai signal (magenta) points to apoptotic bodies located basally in the epithelium. a: apical. Grey: E-Cadherin. Inset scale bar: 5 μm. **E:** Segmentation output from the disc in **D**, showing the DSRF pattern and the GC3Ai segmented signal (magenta). Veins regions are grey. Black dots are landmarks. **F:** Superimposition of the segmentation data from 29 discs scaled, rotated and aligned using the General Procrustes Analysis (GPA), showing a spatially heterogeneous signal for GC3Ai (magenta). The map shown for vein / intervein territories is the average map from the 29 discs. **G-I:** Heat-maps showing the average GC3Ai positive area on each of the 40 compartments of the wing disc for the *control* (**G**, n = 29), *Dronc^DN^* (**H**, n = 26) and *hid-RNAi* (**I**, n = 25) genotypes. Numbers within each compartment show the average value for GC3Ai positive area (arbitrary units). These values are also shown as a colour coded heat-map which has the same scale for three maps shown in G-I. **J:** Visualisation of caspase-3 activity using the GC3Ai reporter (magenta) expressed under the control of GMR11F02-GAL4 at ~ 72 h AEL. The wing disc is divided into four quadrants (Anterior-Dorsal; Anterior-Ventral; Posterior-Dorsal; Posterior-Ventral) using the immunostaining anti Wingless (wg, green) and anti Patched (ptc, green) marking compartment boundaries. **K:** Segmentation of the disc shown in **J**. The circled black crosses denote the centroids of the segmented GC3Ai signal. The black line was manually drawn using anti-Wingless and anti-Patched immunostainings to delineate the four quadrants, which are coloured. To account for uncertainty in centroid location of the GC3Ai spots and of manual drawing of the contours, an overlap between the quadrants was allowed. **L:** Mean GC3Ai positive area for 10 discs at ~ 72 h AEL. One red dot represents one disc, blue circles are means and error bars are 95 % confidence intervals. The inset shows the p-values for all pairwise t-tests between quadrants. **M:** Pattern of *hid*^ΔGFP^ (KI) on a single wing disc (LUT fire, Gaussian blur 0.75). White lines denote the vein/intervein territories extracted from the anti-DSRF immunostaining (not shown). **N:** Heat-map showing the mean *hid^ΔGFP^* signal on each of the 40 compartments over n = 17 discs. To allow comparison of intensities among discs, the GFP signal intensity was normalised within each disc (i.e., within each disc, the value of each pixel was divided by the mean signal intensity over all the pixels of the disc). All scale bars are 50 μm length except in **B**, 500 μm, and in **J**, 25 μm.

Accordingly, apoptosis was proposed to buffer developmental fluctuations to ensure reproducible wing size [18]. This was assumed to be based on cell competition: the context dependent elimination of suboptimal cells from growing tissues[19]. Interestingly, while the studies of cell competition are mostly based on the analysis of mutant clone disappearance in the wing disc, we know virtually nothing about spontaneous clone disappearance during normal wing development and the prevalence, localisation and functions of physiological cell competition.

Here, we performed systematic mapping of apoptosis using a live marker of caspases and spatial registration in the larval wing imaginal disc. Unexpectedly, we found striking reproducible biases in the distribution of cell death which outlined hot-spots of apoptosis. These hot-spots correlate with a local increase of clone disappearance probability and significantly reduce local net-growth in the wing disc. Using morphometric analysis, we further demonstrate that these hot-spots can tune adult wing size and shape. Altogether, we reveal that apoptosis cannot be neglected for the study of growth/size and is also an essential modulator of local shape and growth even in a fast-expanding tissue. Our study also proposes a roadmap for a more systematic characterisation of the contribution of cell death to clone dynamics and tissue growth.

## Results

### Caspase activity is spatially biased in the larval wing pouch

We first systematically evaluated the distribution of apoptotic cells in the *Drosophila* wing disc at the larval wandering stage (96 h After Egg Laying - AEL). We used the live effector caspase reporter GC3Ai [20] driven by a wing specific driver (GMR11F02-GAL4, **Figure 1C**). The brightest GFP spots revealed by the reporter correspond to cell debris located on the basal sides, thus staining cells which already extruded ([20], **Figure 1D**). We confirmed the accuracy of the GC3Ai marker as we obtained a similar pattern with cleaved caspase3 staining (cleaved DCP1), although this was less sensitive and fails to mark the cells with faint GC3Ai signal (**Figure 1suppE-G**). To obtain an averaged map of the spatial distribution of cell death, we used spatial landmarks, whose positions were defined by the intervein marker Drosophila Serum Response Factor (DSRF, **Figure 1D,E**). The landmarks positions were used to divide every dissected wing disc into 40 sub-compartments (**Figure 1suppA-D**). In addition, we applied procrustes transformation (translation, rotation and scaling) on landmarks positions to align the discs and superimpose GC3Ai spatial data from many individuals on a single image (see **Methods**). The averaged distribution of apoptotic cells revealed a striking non-homogenous distribution of apoptosis with the strongest “hot spot” located in the anterior - dorsal quadrant, near the dorsal-ventral boundary (**Figure 1 F,G**). This bias is not driven by the driver or any side effect of GC3Ai as a similar pattern was obtained in the *w*^-^ background with cleaved caspase3 staining (**Figure 1suppH-J**), and with GC3Ai driven by another GAL4 (*nubbin-gal4*, **Figure 1suppK**). To check whether the spatial bias of GC3Ai signal persists during wing disc development, we also monitored GC3Ai-positive cells in the early L3 wing disc (72h AEL). As we could not use the same spatial landmarks (the intervein regions are not yet defined at this stage) we instead used the location of the 4 compartments defined by AP and DV compartment boundaries (Anterior, Posterior, Dorsal and Ventral) as a spatial reference. The number of apoptotic cells was more than two times higher in the anterior-dorsal compartment than in the others (**Figure 1J-L**). This suggested that the spatial bias emerges at least during early L3 stage and persists for more than 24 hours. We then checked whether this pattern was indeed dependent on core regulators of apoptosis/caspases. Co-expression of GC3Ai with a dominant negative allele of Dronc *(Drosophila* caspase9) significantly reduced the amount of GC3Ai signal and reduced the spatial bias (**Figure 1H**, **Figure 1suppL-N, R**). Similarly, depletion of the pro-apoptotic gene *hid* by RNAi in the wing pouch almost completely abolished the GC3Ai signal (**Figure 1I**, **Figure 1suppO-Q, R**). This suggests that the spatial bias is indeed caspase-dependent and most likely relies on the expression of the pro-apoptotic gene *hid*. Accordingly, we found a slight but systematic and reproducible increase of *hid* expression near the GC3Ai hot spot region using a GFP insertion at the *hid* locus (**Figure 1M,N**). Note however that other factors may also contribute to the spatial bias as we still observed some heterogeneity in cell death distribution upon Dronc inhibition or *hid* depletion (**Figure 1suppN, Q, S**).

Altogether, we conclude that contrary to what was previously observed [17], apoptosis is not spatially homogeneous in the wing imaginal disc and that some regions (anterior and close to the DV boundary, dorsal compartment) undergo higher rates of apoptosis. These hot-spots are in part explained by a spatial bias in the expression of the pro-apoptotic gene *hid*.

### The spatial bias in caspase activity influences clonal disappearance and local net-growth

We next checked whether this spatial bias in apoptosis could have any consequence on clonal dynamics and local growth. Since ex-vivo live imaging of the wing disc does not allow long-term tracking of cell fate and local growth, we used an alternative approach based on twin-clone labelling. We first used the QMARCM technique [21] to stain the two daughter cells generated after mitotic recombination and their progeny with different fluorescent markers using alternate sets of transcription factors (Gal4 or the Qsystem, **Figure 2A,B**). Since cell movements and cell-cell intercalations are relatively neglectable in the wing disc [22, 23], the spatial proximity of patches of cells marked with GFP and RFP can be unambiguously attributed to a twin-clone in conditions of low frequency of clone induction. We assessed the probability of clone disappearance by scoring the number of single-coloured clones which are not in the vicinity of the other sibling clone 48h after clone induction. Such an observation can only be explained by the early disappearance of the lineage of the other daughter cell, and thus identifies a clone disappearance event (**Figure 2A**). We first noted that the QMARCM system has an intrinsic bias towards smaller clones and higher probability of clone disappearance for GFP/Gal4 clones relative to RFP/QF ones (**Figure 2suppA**). This however should not preclude the analysis of potential spatial biases. By multiplexing a large number of discs and using the same spatial landmarks as those used in **Figure 1suppC** (**Figure 2B**), we obtained a coarse-grained spatial map of the probability of clone disappearance (**Figure 2C**). Strikingly, we observed that similar to caspase activation in the tissue, the probability of clone disappearance is not spatially homogenous. We identified clear hot-spots of clone death, including the anterior-dorsal region near the DV boundary (~two-fold increase compared to posterior compartments at similar DV positions, **Figure 2C, 2suppB**), as already outlined by the caspase positive cells mapping (**Figure 1G**). Note that this clone strategy reflects the pattern of apoptosis at earlier stages of development compared to the one revealed with GC3Ai. A similar pattern was observed using an alternative twin-clone method (twin-spot MARCM [24], using RNAi depletion of RFP and GFP via a GAL4 expressed in the wing pouch, **Figure 2D, 2suppB**) suggesting that the pattern of clone disappearance is not specific to the clone induction system. Importantly, clone size was also on average smaller (estimated through total apical surface and number of cells) in regions showing high rates of apoptosis and clone disappearance (**Figure 3A, Figure 3suppC,E**). This suggested that spatial biases in caspase activity not only modulate the probability of clone survival, but also the cumulative local growth in the wing disc. Indeed, these local differences in clone size match the differences numerically estimated from an exponential growth model with no spatial difference in proliferation rate and including the spatial differences in apoptotic rates calculated from our measurements of clone disappearance (see **Methods, Figure 3B-D**). This indicates that local differences in clone size could be solely explained by spatial differences in apoptotic rates. To check whether these spatial differences are indeed driven by caspases, we repeated the QMARCM clonal assay upon inhibition of caspase in the Gal4 sibling clones (using UAS-Dronc^DN^). While this did not totally abolish clone disappearance, the spatial pattern was flattened with no visible hot-spot of clone disappearance in the anterior side (**Figure 2C’, 2suppB**). The spatial pattern of clone disappearance was also affected upon global inhibition of *hid* by RNAi with the twin-spot MARCM assay (**Figure 2D’, 2suppB**). These two approaches also lead to more homogenous clone size between anterior and posterior compartments in the wing disc (**Figure 3suppD,F**).

**Figure 2:**
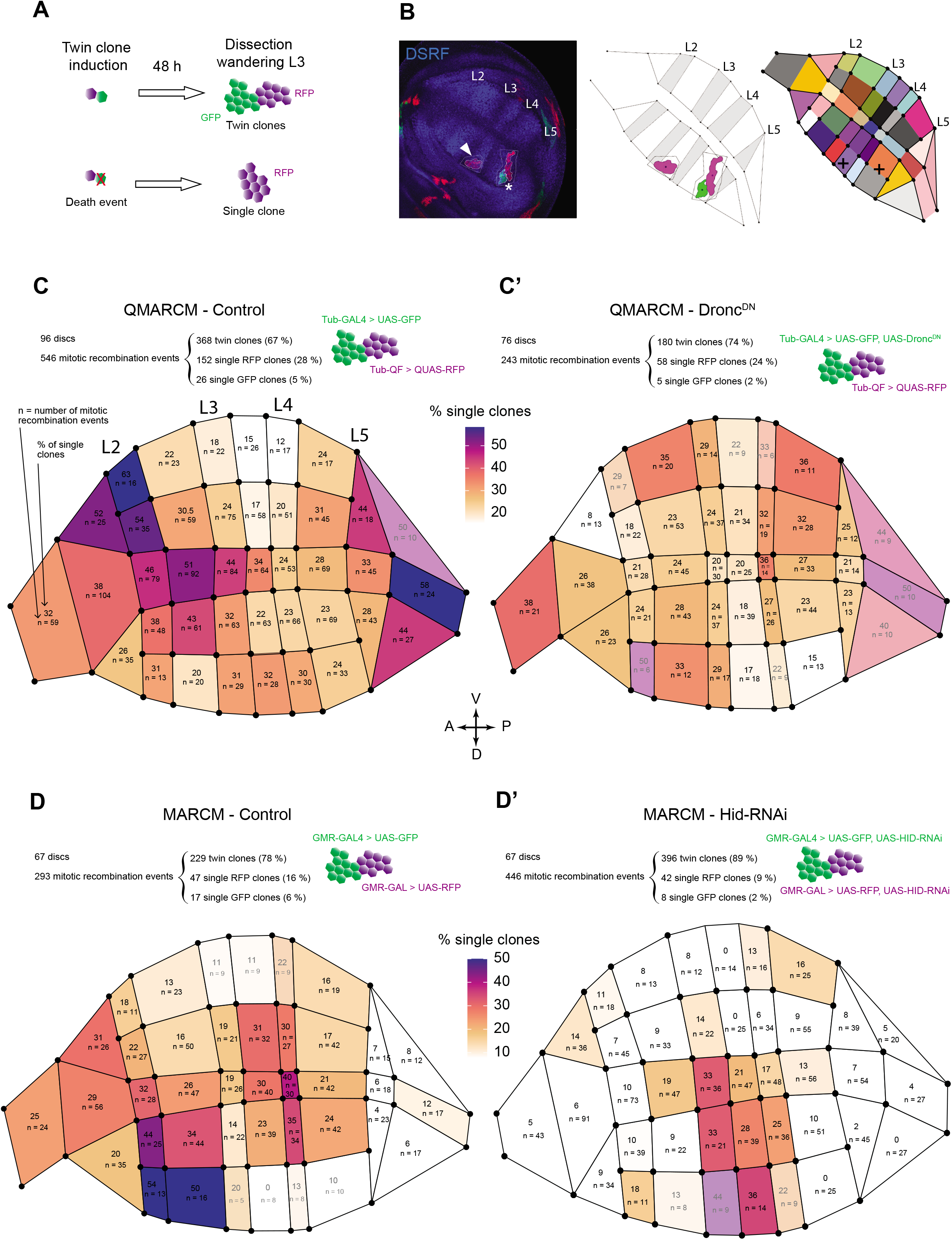
clone disappearance is spatially patterned in the developing wing. **A:** Rationale for inferring cell death events from twin-clone induction experiments. Upon twin-clone induction and 48 h of growth, the observation of twin clones composed of one GFP and one RFP clone is expected. However, early occurrence of cell death events after clone induction may eliminate one sibbling resulting in the occurrence of a single GFP or RFP clone without any twin counterpart in its vicinity. **B:** Spatial mapping for twin and single clone occurrences in the wing disc. Left, image of a wing disc at 96 h AEL stained with anti-DSRF and showing a single RFP clone (arrowhead) and a twin clone GFP-RFP (asterisk). Middle, manual segmentation of the tissue using the DSRF signal, and automated segmentation of the clones. Right, spatial map of the wing disc divided in 40 compartments (following the DSRF-based segmentation) into which clones positions are mapped (black crosses show location of clones centroids). **C-D:** Spatial maps showing the pattern of single clone occurrences under different genetic conditions. In each case, the genotypes of the clones, the number of discs studied, the total number of mitotic recombination events, of twin clones, and of single clones observed are given. Colour scale of the heat maps shows the percentage of single clones observed in each compartment, relative to the total number of mitotic recombination events assigned to this compartment (n). For clarity, the compartments for which n ≤ 10 were shaded, since the inferred proportions from such low sample size are very unreliable.**C-C’:** Spatial map of single clone occurrences obtained by inducing twin clones using the twin spot QMARCM system, in control conditions (**C**) and by additionally expressing UAS-Dronc^DN^ in the GFP clone (**C’**). **C** and **C’** share the same colour scale. **D-D’:** Spatial map of single clone occurrences obtained by inducing twin clones using the twin spot MARCM system driven by the GMR11F02-GAL4, in control conditions (**D**) and by additionally expressing *UAS-hid-dsRNA* in all the pouch (**D’**). D and D’ share the same colour scale.

**Figure 3.**
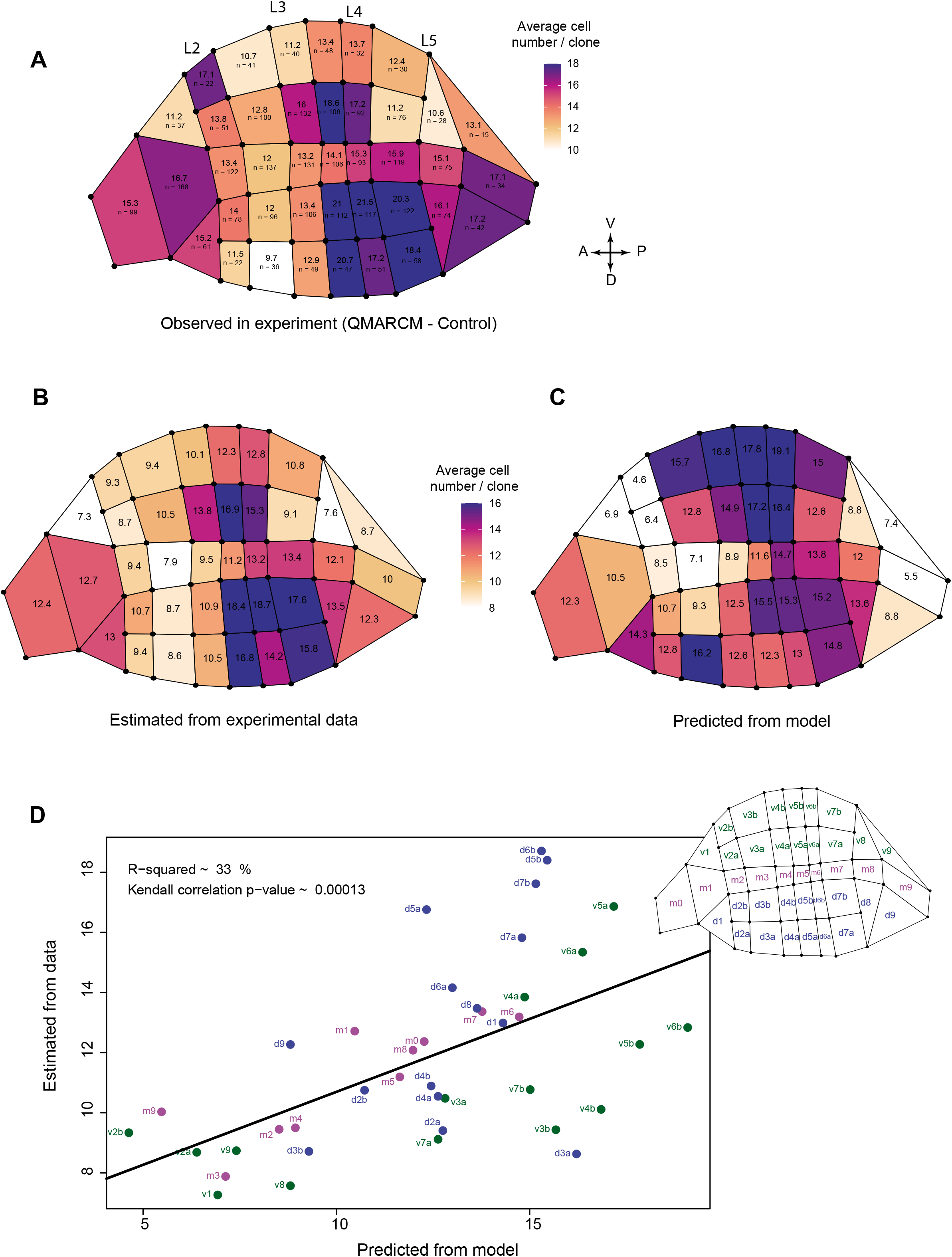
Local variation of clone size can be predicted by the local variation of apoptosis. **A:** Spatial map for clone size (expressed in mean number of cells/clone) in QMARCM control experiment. Clones were considered irrespective of their colour. For each compartment, the average clone surface was calculated and divided a posteriori by the average cell apical size obtained in the same compartment to obtain an estimation of the averaged cell number per clone (see **Methods** and **Figure 3supp**). **B:** Spatial map of the mean value of clone size (μ(*t*), see **Methods**) estimated by integrating the experimental average clone size for each compartment as well as the “0 values” (estimated from the proportion of singe colour clone, see **Methods**). **C**: Spatial map of the averaged predicted number of cells per clone assuming a constant and homogenous growth rate throughout the pouch and heterogeneous rate of apoptosis estimated from the local proportion of single coloured clone (see **Methods**). B and C share the same colour scale bar. **D:** Correlation between the average cell number per clone predicted from the model and estimated from experimental data for each compartment. The black line is the linear regression. The R-squared of the regression and the p-value of a correlation test (Kendall) are shown. Each dot is the data for one compartment, coloured and named according to the map shown on the right side of the plot (blue: dorsal compartments d1 to d9; magenta: margin compartments m0 to m9; green: ventral compartments v1 to v9).

Altogether, we conclude that the spatial bias in *hid* expression and caspase activity generates a hot-spot of clone disappearance which also significantly reduces local growth rate mostly in the anterior-dorsal region near the DV boundary. This suggests that local increases of caspase activity could play an instructive role in significantly modulating local growth, which therefore may affect the final shape and size of the tissue.

### Patterned caspase activity modulates adult wing shape and size

We therefore checked whether local biases in caspase activity could significantly impact adult wing shape and size. To obtain a precise and quantitative description of adult wing shape and size, we used a quantitative assay based on semi-automatic wing segmentation, landmark positioning and procrustes alignment of wings (which includes translation, rotation and global rescaling) [25, 26] (**Figure 4suppA**). Interestingly, inhibition of cell death in the wing tissue using *hid* RNAi led to a significant increase in wing size (**Figure 4B**, 5.5%, a similar range to what was obtained upon ectopic overexpression of Dp110/PI3K [27]), suggesting that apoptosis has a net negative effect on adult tissue size. We also observed significant changes to adult wing shape, including a global increase in the size of the most anterior and posterior domains, as well as a global wing rounding (**Figure 4C,E,F**). To check whether local increases in apoptosis were indeed responsible for the local modulation of shape and size, we used various drivers to inhibit apoptosis in different wing subdomains (**Figure 4A**). Accordingly, inhibition of *hid* in the patched domain (*ptc-gal4*, expressed in a band near the AP compartment boundary, **Figure 4A**) led to a global increase of wing size dominated by the expansion of the region overlapping the *ptc* domain (**Figure 4B,C**). This suggested that apoptosis has a local impact on growth even in regions other than the apoptosis hot-spots identified at larval stage. We also used drivers restricted to the DV boundary either in the anterior (encompassing the caspase hot-spot, **Figure 1G**) or posterior part of the wing disc (**Figure 4A**). Interestingly, while *hid* depletion in the posterior-DV boundary domain had a mild effect on wing size (similar to inhibition of *hid* in the ptc domain), this effect was enhanced upon inhibition in the anterior-DV boundary domain reaching differences close to the one observed upon inhibition of *hid* in the full wing pouch (4.1% versus 5.5% increase, **Figure 4B**). This suggests that apoptosis around the anterior DV margin (the strongest hot-spot of apoptosis) can account for a significant proportion of the effect on adult wing size. Importantly, the expansion of the most anterior domain of the wing was only observed upon inhibition of *hid* in the anterior-DV boundary domain (**Figure 4C**), suggesting again that apoptosis has a local effect on size. The observed effects on adult size might also be due to changes during the pupal stage. Therefore we also performed experiments with conditional *hid* depletion to assess the relative contribution of *hid* expression at larval stage versus pupal stage to the final wing shape (**Figure 4suppB,C**). While depletion of *hid* starting at early-mid pupal stage only had a minor effect on wing shape (**Figure 4D, 4suppE**), *hid* depletion from early larval stage to early pupal stage was sufficient to recapitulate the rounding of the wing and the main shape modulation observed upon *hid* depletion throughout development (**Figure 4D-F, 4suppE**). Note that we could not use this context to analyze the effect on absolute size, as it was completely dominated by the well-documented effect of temperature on wing size [28, 29] (**Figure 4suppD**).

**Figure 4:**
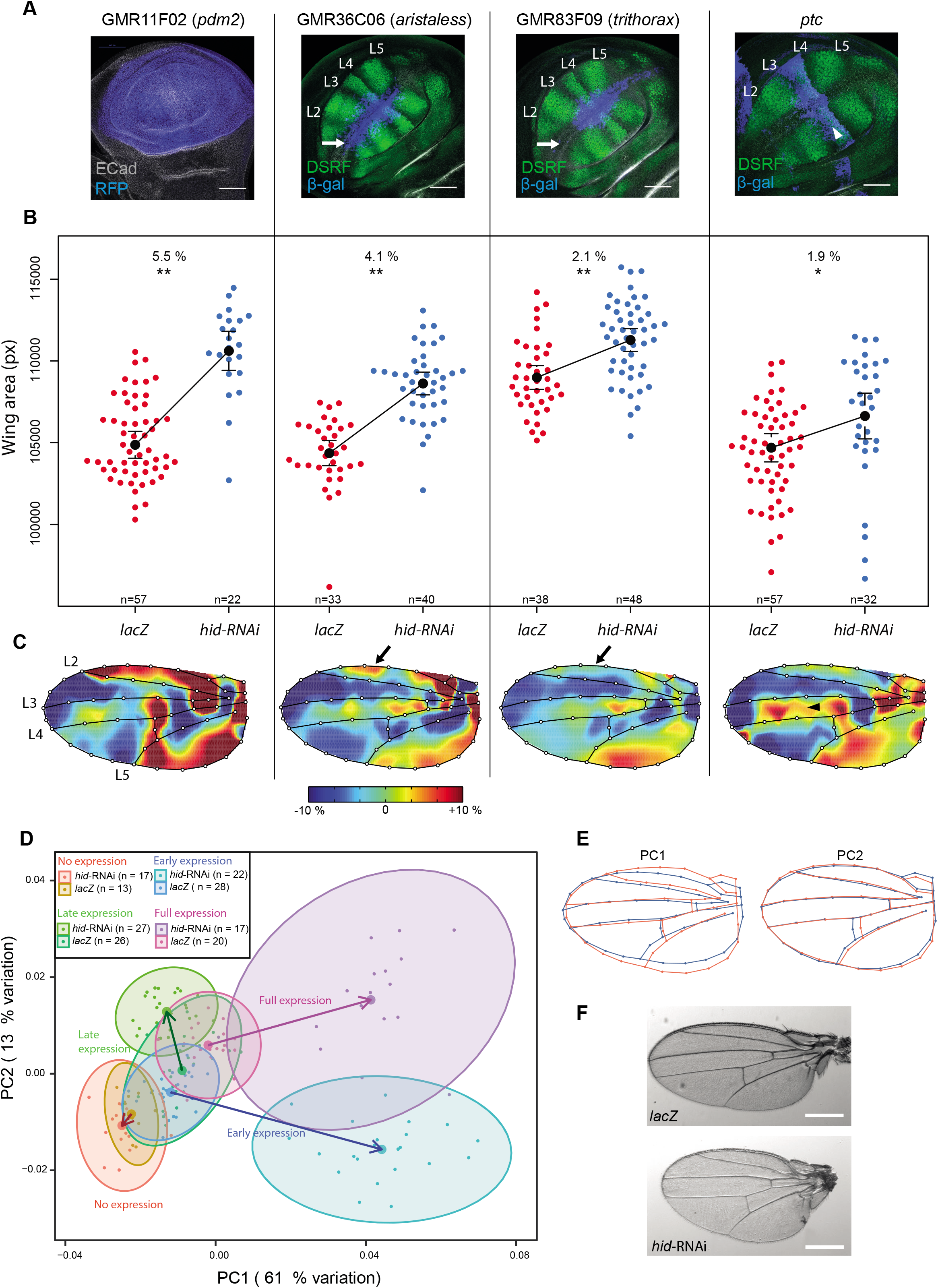
Apoptosis affects locally and globally wing size and shape during early stages of development. **A:** Pattern of expression at ~ 96 h AEL of the GAL4 drivers used to express *hid-RNAi*. White arrows show the most anterior region of the disc, where GAL4 is expressed in the case of *aristaless-GAL4*, but not in the case of *trithorax*-GAL4. White arrowhead highlights the *ptc* domain. **B:** Variation of adult wing size caused by expression of *hid*-RNAi using different GAL4 drivers. For each GAL4 driver shown in **A**, the effect of expressing *hid*-RNAi compared to *lacZ* is shown. Blue and red dots represent individual wing size values, black dots are means, and error bars are 95 % confidence interval of the mean. For each driver, the % of variation of the *hid*-RNAi mean relative to the *lacZ*mean, as well as the results of pairwise t-tests between *lacZ* and *hid*-RNAi groups are shown. **: p < 0.001; *: p < 0.01. « n » denotes the number of wings for each group. **C:** Local variation of tissue shape driven by expression of *hid*-RNAi with the GAL4 drivers shown in A. Colours represent changes in relative area necessary to transform the average wing from the *lacZ* group into the average wing of *hid*-RNAi group (red: increase, blue: decrease, number of wings shown in **B**). Colour bar shows the upper and lower limits in deformation. Highest colour intensity is reached at 10 % increase and decrease in local area in *hid*-RNAi group with respect to *lacZ* group. Arrows / arrowhead show the regions corresponding to the regions pointed by arrows / arrowhead in **A**. **D:** Principal components analysis (PCA) showing variations of wing shape (according to the positions of 49 Procrustes-aligned landmarks and semilandmarks) caused by expression of *hid*-RNAi at different developmental stages (using Gal80^ts^ and switch between 18°C and 29°C). Each small dot represents one wing shape. 8 conditions are shown (no expression, early expression, late expression and full expression of *hid*-RNAi or *lacZ* with GMR11F02-GAL4 driver). Big dots represent the mean of each condition. Vectors connect the mean of the *lacZ* group to the one of *hid*-RNAi group of the corresponding condition, thus allowing to visualise magnitude and direction of the effect on wing shape due to *hid*-RNAi expression. Ellipses are 95 % confidence intervals of the means. **E:** Diagrams showing the variation of wing shape along the first and the second principal component (PC) axes of the PCA shown in **D**, from low PC values (blue) to high values (red). **F:** Pictures of mounted wings illustrating the shape rounding observed upon *hid*-RNAi expression during early stages as compared to expression of *lacZ*. Scale bars are 500 μm.

To have a more quantitative description of shape changes induced by these genetic backgrounds, we used Principal Components Analysis (PCA) applied to the 48 landmarks defined by the wing segmentation (**Figure 4suppA**). The combinatorial effect of *hid* depletion during larval and pupal stages is well reflected by this PCA analysis (**Figure 4D**), which can decompose the total effect of *hid* depletion throughout development (“full expression”, purple arrow) into a strong effect coming from the larval stage along Principal Component 1 (“early expression”, blue arrow, along PC1) and a mild effect during pupal stage (“late expression”, green arrow, mild effect along PC2). Accordingly, PC1 mostly reflects the rounding of the wing (**Figure 4E**, accounting for 61% of the variation in the PCA analysis) while PC2 mostly relies on the shift of the posterior cross-vein (**Figure 4E**, accounting for 13% of the variation of the PCA analysis). The effect of this temporal depletion, combined with the wing pouch specificity of the other drivers strongly argue for the important role of *hid* expression pattern in the pouch during larval development on adult wing shape.

Altogether, we conclude that spatial biases in apoptosis distribution in the growing wing disc can significantly modify the shape of the adult wing by locally reducing net growth. This suggests that the fine spatial tuning of apoptosis in a fast-growing tissue plays an instructive role for organ shape and size regulation.

## Discussion

In this study, we outlined an unexpected pattern of apoptosis in the growing wing imaginal disc with a significant upregulation in the anterior and dorsal compartment of the wing. To our knowledge this is the first demonstration that apoptosis is spatially biased in the wing imaginal disc. Moreover, our quantitative assessments of clonal growth and adult wing shape clearly show that this local upregulation of apoptosis has a significant impact on local net growth and final adult shape. So far, the characterisation of cell death has never been performed in a systematic manner using spatial landmarks allowing superimposition of data from many individuals, which may explain why such biases have been missed [17]. Similarly, the impact of apoptosis on tissue shape and size has never been studied with such quantitative readouts. More generally, the assessment of apoptosis levels and distributions is usually based on cleaved caspase3 and TUNEL staining [11]. However, in conditions where the lifetime of apoptotic debris is relatively short, these assays may easily miss potential patterns of interest and may underestimate the real contribution of apoptosis. Our work emphasises the need for more systematic quantitative assessments of death distribution in order to reach solid conclusions about the contribution of apoptosis to organ shape/size regulation. The pipeline used in this study may easily be applied in other developmental contexts which are not amenable for long term live imaging, as long as markers can be used for spatial registration and tools are available for clone generation.

Compensatory proliferation is one of the best studied processes that relates apoptosis to the induction of cell proliferation [30]. So far, this process was mostly characterised either in conditions of massive death induction through irradiation and genetic induction of apoptosis in large domains [31, 32], or through the perturbation of the core apoptotic pathway (e.g.: by blocking some of the essential caspases, caspase3 in *Drosophila*[33], or caspase9 in mammalian epidermis[34]). However, to our knowledge there is no study that has clearly identified biases in cell proliferation distribution in the vicinity of physiological apoptosis *in vivo*. Surprisingly, we observed a net negative effect of the local increase of cell death on local growth and the size of the final adult compartment, and also outlined a significant effect of physiological death on the final size of the adult wing (**Figure 2** and **Figure 4**). This suggests that compensatory proliferation is unlikely to occur in the context of physiological apoptosis, or at least that its contribution is relatively minor and not sufficient to compensate for cell loss by apoptosis. Further quantitative studies of the coupling between apoptosis and cell proliferation will be essential to assess its real contribution to physiological growth regulation. Interestingly, a recent study outlined a significant positive upregulation of proliferation near dying MDCK cells, however this effect is strongly context-dependent and is not visible at low stiffness or high density values[35], conditions that may apply to the wing imaginal disc.

Cell competition is the process describing the context-dependent elimination of viable but suboptimal cells [6]. Studies of clonal elimination in the wing imaginal disc have thoroughly contributed to our understanding of this process. Interestingly, most of these studies assumed that clone disappearance in physiological conditions is largely neglectable. Our results suggest that a significant proportion of WT clones may undergo early elimination in the wing imaginal disc, and that this elimination is spatially biased. On the one hand, this opens the possibility for spontaneous competition occurring in the wing imaginal disc, which so far has not been thoroughly explored (although its existence has been suggested by indirect genetic evidence [7, 18, 36]). On the other hand, it also suggests that cell elimination during competition may be influenced by this spatial pattern of caspase activity and apoptosis. Further quantitative description of clonal elimination during competition may reveal such spatial bias and would help to study the influence of pre-existing patterns of caspases/apoptosis on cell competition and tissue plasticity.

In this study, we showed that local modulation of *hid* expression can fine-tune the local level of apoptosis which will impact local growth and adult tissue shape in a subtle and quantitative way, similarly to the phenotypic changes observed at the macroevolutionary scale [37]. As such, the evolution of the cis-regulatory elements of pro-apoptotic genes may constitute an additional lever for shape evolution that could be used to fine tune adult appendage shape. Interestingly, these pro-apoptotic genes are most likely less pleiotropic than pro-proliferative pathways such as morphogens, RTK signaling pathways, or Hippo pathways that are classically studied for the regulation of size and growth [10]. As such, modulating pro-apoptotic genes levels and pattern of expression may constitute a relatively parsimonious way to evolve wing and appendage shape.

## Acknowledgements

We thank members of RL lab for critical reading of the manuscript. We would like to thank Christina Fissoun, Lucia Rodriguez Vazquez, Gaurav Shajepal and Delia Cicciarello for their contribution to adult wing and wing disc analysis during their internship. We are also grateful to Jean-Paul Vincent, Magali Suzanne, David Houle, Seth Blair, the Bloomington Drosophila Stock Center, the Drosophila Genetic Resource Center and the Vienna Drosophila Resource Center, Flybase for sharing essential information, stocks and reagents. We also thank Benoît Aigouy for the Packing Analyser software and Jean-Yves Tinevez and the image analysis platform of Institut Pasteur for the LocalZprojector plugin on Fiji. Work in RL lab is supported by the Institut Pasteur (G5 starting package), the ERC starting grant CoSpaDD (Competition for Space in Development and Disease, grant number 758457), the Cercle FSER and the CNRS (UMR 3738).

## Authors contribution

RL and AMV discussed and designed the project and wrote the manuscript. TC performed part of the experiments on adult wing shape and twin clones analysis as well as the wing disc segmentation. AD provided the theoretical calculation of the expected distribution of clone size based on differences in apoptotic rates. AMV performed all the other experiments and analysis. Every author has commented and edited the manuscript.

## Methods

### Resource availability

#### Lead contact

Further information and requests for resources and reagents should be directed to and will be fulfilled by the lead contact, Romain Levayer.

#### Material availability

All the reagents generated in this study will be shared upon request to the lead contact without any restrictions.

#### Data and Code availability

All code generated in this study and the raw data corresponding to each figure panel (including images) can be shared upon request and will be uploaded soon to a repository.

### Experimental model and subject details

#### Drosophila melanogaster husbandry

All the experiments were performed using *Drosophila melanogaster* fly lines (listed in **Table 1**) breed with regular husbandry techniques. The fly food used contains agar agar (7.6 g/l), saccharose (53 g/l) dry yeast (48 g/l), maize flour (38.4 g/l), propionic acid (3.8 ml/l), Nipagin 10% (23.9 ml/l) all mixed in one liter of distilled water. Flies were raised at 25°C in plastic vials with a 12h/12h dark light cycle at 60% of moisture unless specified in the legends and in the table below (alternatively raised at 18°C or 29°C). Females and males were used without distinction for all the experiments, except for the adult wing shape analysis in which only left wings from female flies were used. We did not determine the health/immune status of pupae, adults, embryos and larvae, they were not involved in previous procedures, and they were all drug and test naïve.

**Table 1:** description and origin of the *Drosophila melanogaster* strains used in this study.

| Fly line | Chromosome location | Origin (citation) | RRID |
| --- | --- | --- | --- |
| *UAS-GC3Ai /TM6b* | III | [20], Magali Suzanne | none |
| *w; UAS-GC3Ai/Cyo ; MKRS/TM6b* | II | Bloomington and [20] | BDCS_84346 |
| *UAS-hid dsRNA* | III | VDRC | GD 8269 |
| *UAS-lacZ* | II | Bloomington | BDCS_8529 |
| *UAS-dronc<sup>DN</sup>* | II | Boomington | BDCS_58992 |
| *GMR11F02-gal4* | III | Bloomington | BDCS_48928 |
| *w<sup>118</sup>* | X | Bloomington | BDCS_3605 |
| *UAS-dicer2 ; nubbin-gal4* | X, II | Bloomington | BDCS_25754 |
| *w; ; hid<sup>Δ.GFP</sup>/TM3* | III | [38], Jean-Paul Vincent | none |
| *yw; ET40-QF, QUAS-tdTomato; FRT82B, tub-QS* | X, II, III | Bloomington | BDCS_30042 |
| *yw, hs-flp22, UAS-GFP ; if/Cyo ; tub-gal4, FRT82B, tub-gal80/TM6b* | X, III | This study and<br>Bloomington | BDCS_86311 |
| *yw; UAS-mCd2RFP , UAS-dsRNACD8*<br>*FRT40A ; TM3/TM6b* | II, III | Bloomington | BDCS_56184 |
| *yw; UAS-mcd8GFP, UAS-dsRNACD2*<br>*FRT40A ; TM3/TM6b* | II, III | Bloomington | BDCS_56185 |
| *w; ptc-gal4* | II | Bloomington | BDCS_2017 |
| *w; GMR83F09-gal4* | II | Bloomington | BDCS_40367 |
| *w; GMR36C06-gal4* | II | Bloomington | BDCS_49931 |
| *w; tub-gal80ts* | II | Bloomington | BDCS_7108 |
| *w; UAS-lacZ* | II | Bloomington | BDCS_8529 |

#### Drosophila melanogaster strains

The strains used in this study and their origin are listed in the table 1 below.

The exact genotype used for each experiment is listed in table 2

**Table 2:**
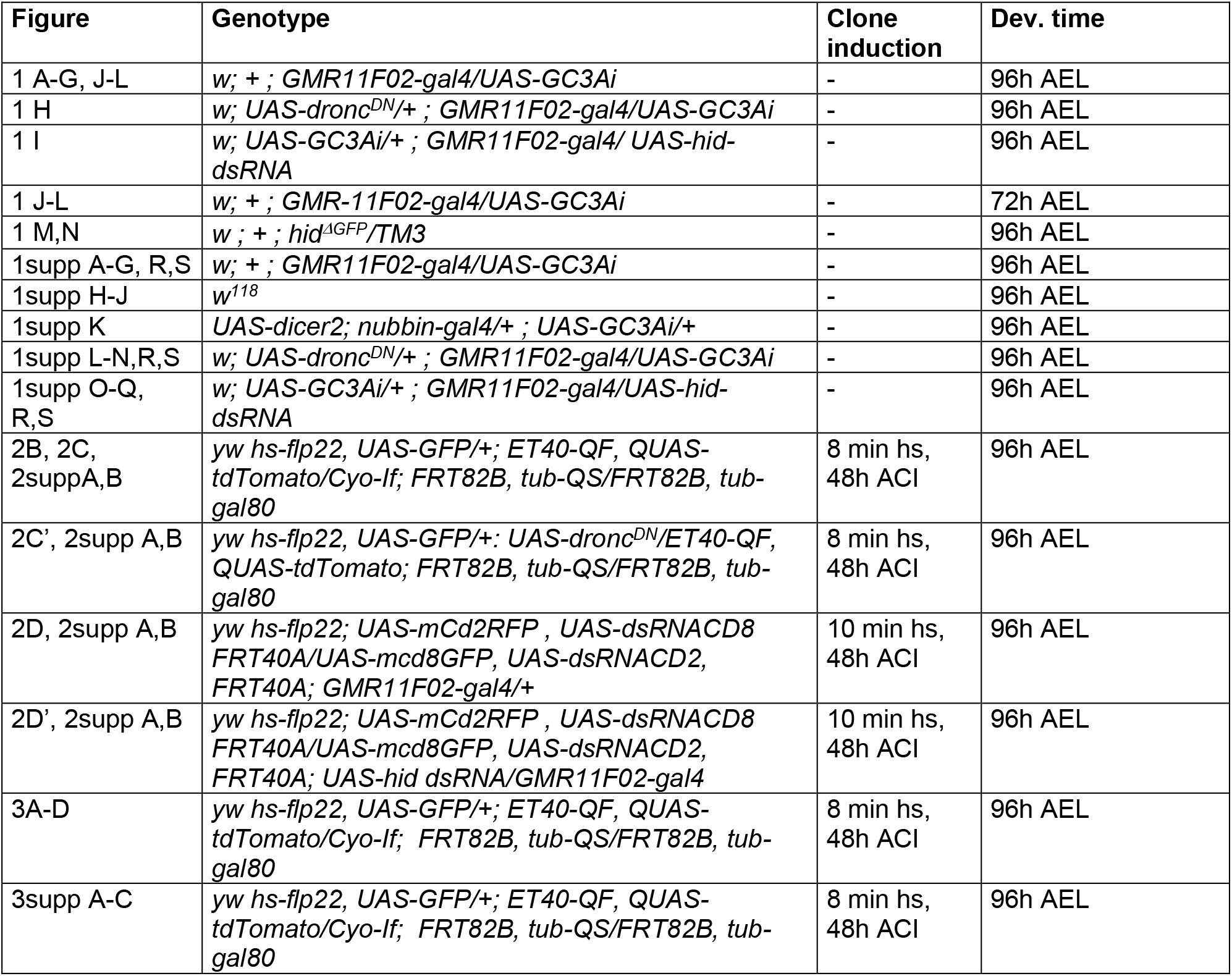
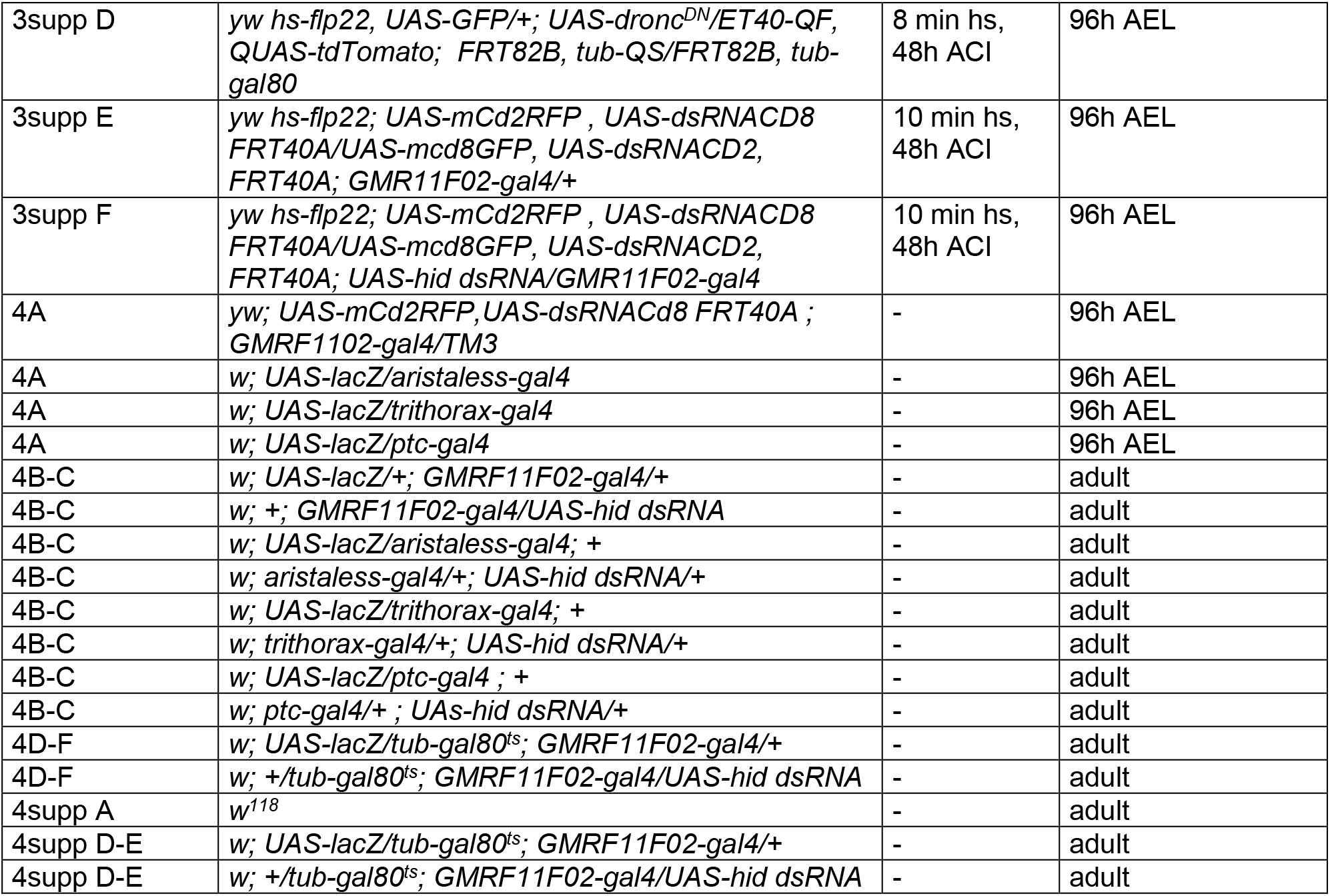
genotype used for each experiment. AEL: after egg laying; hs: heat shock duration at 37°C; ACI: time after clone induction.

| Figure | Genotype | Clone<br>induction | Dev. time |
| --- | --- | --- | --- |
| 1 A-G, J-L | *w; + ; GMR11F02-gal4/UAS-GC3Ai* | - | 96h AEL |
| 1 H | *w; UAS-dronc<sup>DN</sup>/+ ; GMR11F02-gal4/UAS-GC3Ai* | - | 96h AEL |
| 1 I | *w; UAS-GC3Ai/+ ; GMR11F02-gal4/ UAS-hid-dsRNA* | - | 96h AEL |
| 1 J-L | *w; + ; GMR-11F02-gal4/UAS-GC3Ai* | - | 72h AEL |
| 1 M,N | *w ; + ; hid<sup>ΔGFP</sup>/TM3* | - | 96h AEL |
| 1supp A-G, R,S | *w; + ; GMR11F02-gal4/UAS-GC3Ai* | - | 96h AEL |
| 1supp H-J | *w<sup>118</sup>* | - | 96h AEL |
| 1supp K | *UAS-dicer2; nubbin-gal4/+ ; UAS-GC3Ai/+* | - | 96h AEL |
| 1supp L-N,R,S | *w; UAS-dronc<sup>DN</sup>/+ ; GMR11F02-gal4/UAS-GC3Ai* | - | 96h AEL |
| 1supp O-Q, R,S | *w; UAS-GC3Ai/+ ; GMR11F02-gal4/UAS-hid-dsRNA* | - | 96h AEL |
| 2B, 2C, 2supp A,B | *yw hs-flp22, UAS-GFP/+; ET40-QF, QUAS-tdTomato/Cyo-If; FRT82B, tub-QS/FRT82B, tub-gal80* | 8 min hs, 48h ACI | 96h AEL |
| 2C', 2supp A,B | *yw hs-flp22, UAS-GFP/+; UAS-dronc<sup>DN</sup>/ET40-QF, QUAS-tdTomato; FRT82B, tub-QS/FRT82B, tub-gal80* | 8 min hs, 48h ACI | 96h AEL |
| 2D, 2supp A,B | *yw hs-flp22; UAS-mCd2RFP , UAS-dsRNACD8 FRT40A/UAS-mcd8GFP, UAS-dsRNACD2, FRT40A; GMR11F02-gal4/+* | 10 min hs, 48h ACI | 96h AEL |
| 2D', 2supp A,B | *yw hs-flp22; UAS-mCd2RFP , UAS-dsRNACD8 FRT40A/UAS-mcd8GFP, UAS-dsRNACD2, FRT40A; UAS-hid dsRNA/GMR11F02-gal4* | 10 min hs, 48h ACI | 96h AEL |
| 3A-D | *yw hs-flp22, UAS-GFP/+; ET40-QF, QUAS-tdTomato/Cyo-If; FRT82B, tub-QS/FRT82B, tub-gal80* | 8 min hs, 48h ACI | 96h AEL |
| 3supp A-C | *yw hs-flp22, UAS-GFP/+; ET40-QF, QUAS-tdTomato/Cyo-If; FRT82B, tub-QS/FRT82B, tub-gal80* | 8 min hs, 48h ACI | 96h AEL |
| 3supp D | *yw hs-flp22, UAS-GFP/+; UAS-dronc<sup>DN</sup>/ET40-QF, QUAS-tdTomato; FRT82B, tub-QS/FRT82B, tub-gal80* | 8 min hs, 48h ACI | 96h AEL |
| 3supp E | *yw hs-flp22; UAS-mCd2RFP, UAS-dsRNACD8 FRT40A/UAS-mcd8GFP, UAS-dsRNACD2, FRT40A; GMR11F02-gal4/+* | 10 min hs, 48h ACI | 96h AEL |
| 3supp F | *yw hs-flp22; UAS-mCd2RFP, UAS-dsRNACD8 FRT40A/UAS-mcd8GFP, UAS-dsRNACD2, FRT40A; UAS-hid dsRNA/GMR11F02-gal4* | 10 min hs, 48h ACI | 96h AEL |
| 4A | *yw; UAS-mCd2RFP, UAS-dsRNACd8 FRT40A ; GMR1102-gal4/TM3* | - | 96h AEL |
| 4A | *w; UAS-lacZ/aristaless-gal4* | - | 96h AEL |
| 4A | *w; UAS-lacZ/trithorax-gal4* | - | 96h AEL |
| 4A | *w; UAS-lacZ/ptc-gal4* | - | 96h AEL |
| 4B-C | *w; UAS-lacZ/+; GMR11F02-gal4/+* | - | adult |
| 4B-C | *w; +; GMR11F02-gal4/UAS-hid dsRNA* | - | adult |
| 4B-C | *w; UAS-lacZ/aristaless-gal4; +* | - | adult |
| 4B-C | *w; aristaless-gal4/+; UAS-hid dsRNA/+* | - | adult |
| 4B-C | *w; UAS-lacZ/trithorax-gal4; +* | - | adult |
| 4B-C | *w; trithorax-gal4/+; UAS-hid dsRNA/+* | - | adult |
| 4B-C | *w; UAS-lacZ/ptc-gal4 ; +* | - | adult |
| 4B-C | *w; ptc-gal4/+ ; UAs-hid dsRNA/+* | - | adult |
| 4D-F | *w; UAS-lacZ/tub-gal80<sup>ts</sup>; GMR11F02-gal4/+* | - | adult |
| 4D-F | *w; +/tub-gal80<sup>ts</sup>; GMR11F02-gal4/UAS-hid dsRNA* | - | adult |
| 4supp A | *w<sup>118</sup>* | - | adult |
| 4supp D-E | *w; UAS-lacZ/tub-gal80<sup>ts</sup>; GMR11F02-gal4/+* | - | adult |
| 4supp D-E | *w; +/tub-gal80<sup>ts</sup>; GMR11F02-gal4/UAS-hid dsRNA* | - | adult |

### Immunostaining

Dissections of larval wing imaginal discs were performed on PBS in ice. Dissected discs were fixed for 20 min in 4 % formaldehyde (SIGMA F8775), rinsed 3 times in PBT (PBS 0.4 % Triton), followed by 10 min permeabilisation in PBT. Primary and secondary antibodies were incubated for 2 h at room temperature (or 12 h at 4 °C) under rocking agitation. After each antibody incubation, discs were rinsed 3 times in PBT, followed by 3 washes of 30 min. Discs were mounted in Vectashield®(EUROBIO SCIENTIFIC / H-1000) and imaged using a confocal spinning disc microscope (Gataca systems) with a 40X oil objective or a LSM880 equipped with a fast Airyscan using a 40X oil objective. The following primary antibodies were used: rat anti E-cadherin (1/100, DCAD2 concentrated DSHB), mouse anti DSRF (1/500, gift of Seth Blair), rat anti Delta (1/1000, gift of François Schweisguth), chicken anti GFP (1/1000, abcam ab13970), rabbit anti RFP (1/500,), rabbit anti DCP-1 (1/100, Cell Signaling 9578S), chicken anti Beta-gal (1/1000, Abcam ab 9361), mouse anti Wingless (1/250, 4D4 concentrated DSHB), mouse anti Patched (1/250, Apa-1 concentrated, DSHB), phalloidin alexa 647 (1/50, Invitrogen). Secondary antibodies were: anti rabbit alexa 555 (1/500, Invitrogen), anti chicken alexa 488 (1/500, Invitrogen), ultrapurified anti mouse 405 (1/500, Jackson ImmunoResearch / 715-476-151), ultrapurified anti mouse CY3 (1/500, Jackson ImmunoResearch / 715-165-151), ultrapurified anti rat 647 (1/500, Jackson ImmunoResearch / 712-605-153).

### Image processing

All images were processed using FIJI[39]. For the twin clones analyses, Z projections of z-stacks were done using the Fiji LocalZProjector plugin using E-cad or phalloidin staining as a reference plane[40], allowing to project a limited number of planes around the apical junction plane while following the local disc curvature. *hid*^ΔGFP^ pattern measurements were done after fixation and GFP immunostaining and maximum z projection. The mean GFP intensity was measured in each compartment (defined with DSRF staining) and normalised by the mean GFP intensity of the full wing pouch.

### QMARCM and twin spot MARCM experiments

Wandering larvae were collected 48 hours after clone induction following a 37°C heat-shock of 8 or 10 minutes (Table 2). Analysis of twin clones was performed on local projections of the wing discs after fixation and staining. Only discs with sufficiently sparse distribution of clones were used (to assign twins unambiguously). To extract clone position and size, each mitotic recombination figure (twin spot or single clone) was manually outlined on FIJI and then automatically segmented by applying a Gaussian blur followed by an automated thresholding (Intermode white method). The centroid of each mitotic recombination figure was obtained by summing the centres of each individual patch of cells composing the twin spot, ponderated by their relative area compared to the total area of the twin clone. Thus, the centroid of each mitotic recombination figure was given by the following formula:

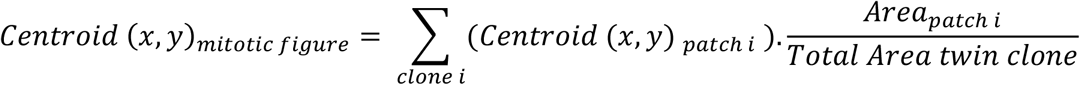

Averaged clone surface was estimated through the segmented surface of GFP or RFP clones (using the local projection around the apical plane of cells). Estimation of the clone size in cell number was obtained a posteriori by segmenting (using Tissue analyser[41]) 3 wing discs stained with E-cad (Figure 3suppA’) and DSRF to position landmarks, and by estimating the averaged apical cell area for each compartment. The 3 wing discs segmented for apical cell area shown a very congruent pattern (Figure 3suppA), thus allowing to pool the data from the 3 discs (Figure supp3 B). The average surface of clone for each compartment was then divided by the local averaged cell size to obtain an estimation of cell number per clone and to correct for effects driven by inhomogeneity of cell apical size throughout the disc.

### Spatial maps of wing discs (twin clones and GC3Ai)

Positions of 36 landmarks were manually set on Z projections of wing discs using FIJI. Landmarks were positioned according to the pattern of DSRF revealed by immunostaining, along the veins, margin and folds of the wing pouch (**Figure 1supp A,B**). Two methods were used to characterise a spatial map of the tissue based on landmarks position. For the first one, landmarks were geometrically aligned within each genotype using the General Procrustes Analysis (GPA) [42] as implemented in R geomorph package[43]. GPA translates the set of landmarks of each wing disc to the same origin, scales them to size, and rotates them until the coordinates of corresponding points align as closely as possible. GPA allows to superimpose as closely as possible several wing discs, thus allowing superimposition of the signal from many individuals (**Figure 1F; Figure 1supp,M,P**). For the second mapping method, 14 additional landmarks were added (landmarks 37 to 50) at the centre of segments defined by other landmarks (for example landmark 37 was set as the centre of the segment defined by landmarks 21 and 22, **Figure 1suppC**). The resulting 50 landmarks were thus used to define polygons dividing the wing disc tissue into 40 compartments (**Figure 1suppC**). With this later approach, clones and apoptotic bodies (defined by GC3Ai or cleaved DCP-1 figures) were assigned to a compartment according to the coordinates of the centre of mass of the clone/apoptotic body. In order to account for uncertainty in clone/apoptotic body localisation (e.g., because of clone movement between the time of induction and imaging), as well as for uncertainty in landmark positioning (e.g., because of immunostaining variability or user error), clone assignment included consideration of an error margin. We used the *st_buffer* function implemented in the *sf* package of R software to add an error margin at a distance of 11.8 μm (40 pixels) around compartment margins (**Fig. 1suppD**). As a result, buffered boundaries of neighbour compartments overlap within each other and a given position in the tissue could be assigned to belong to more than one compartment. This results in a smoothing of the spatial map allowing to account for uncertainty in cells and landmarks positions.

To analyse the spatial pattern of GC3Ai in discs at 72 h AEL, the anti DSRF staining could not be used to draw the landmarks because at this stage the DSRF patterning is not yet established. Instead we stained the wing discs for Ptc and Wg to detect AP and DV compartment and subdivide the wing disc in 4 quadrants (Anterior-Dorsal; Anterior-Ventral; Posterior-Dorsal; Posterior-Ventral).

### Adult wing shape and size analyses

Crosses were done by placing 50-100 couples in egg laying cages with juice-agar plates kept at 25 °C. Freshly hatched larva from the agar plates were transferred to *Drosophila* vials with fly food as described above at a density of 30 individuals / vial, and grown at 18 °C, 25 °C or 29 °C until emergence of adults. Flies were collected and stored in ethanol 70 % at room temperature until dissection. Only left wings from females were taken, and mounted dorsal side up on a glass slide in a solution of 1:1 ethanol 80% and lactic acid 90%. Imaging was carried out on a ZEISS Discovery V8 stereomicroscope using a ZEISS Axiocam ICc 5 camera. All wings were imaged in the same orientation and with the same imaging parameters. For the experiments using Gal80ts (**Figures 4** and **4supp**), flies were crossed in cages supplied with agar petri dishes and allowed to lay eggs for ~ 6 h. Freshly hatched larva were transferred to regular vials at a density of 30 individuals / tube, and placed at 18 °C or 29 °C. Once arrived at the fluid pupal stage, individuals where switched temperature (from 18 °C to 29 °C and vice-versa). Upon hatching, flies were collected for wing dissection.

Wing measurements were done using a semi-automated procedure for estimating the positions of 12 landmarks and 37 semi-landmarks along the wing outline and veins (**Figure 4suppA**). This was done using Wings4 software [25, 26] which fits a B-spline model to the wing from which the coordinates of landmarks and semi-landmarks are extracted. Wings4 outputs were examined using CPR software[26] which allows to screen for outliers and to generate a consolidated dataset of landmarks and wing areas.

Landmarks data were geometrically aligned within each experiment using the General Procrustes Analysis (GPA)[42] as implemented in R geomorph package[43]. GPA translates all wing images to the same origin, scales them to unit-centroid size (centroid size is a measure of specimen size computed as the square root of the sum of squared distances of all the landmarks from the specimen’s centroid) and rotates them until the coordinates of corresponding points align as closely as possible. Differences in landmarks coordinates resulting from the GPA represent shape differences between wings. Wing shape of each individual is characterised by the value of 96 variables, coming from the Procrustes transformation of x and y coordinates of the 48 landmarks and semi-landmarks. To reduce dimensionality of the data, wing shape variation was analysed by Principal Components Analysis as implemented in the function *plotTangentSpace* (now deprecated and replaced by *gm.prcomp*) of the R *geomorph* package. Visualisation of the wing shape variation among the principal component axes was done by comparing wing shape of the individual presenting the lowest value along the axis, with the individual presenting the highest value. To enable visualisation of local growth differences in the adult wing upon apoptosis inhibition, we used the program Lory[44] to show one pattern of relative expansion or contraction that can transform mean shape of control genotype into mean shape of genotypes where apoptosis was inhibited.

### Estimation of clone size differences in an exponential growth regime

The aim of this analysis is to predict spatial differences of clone size under an exponential growth regime given the estimated spatial differences of apoptosis rate based on the twin-clone experimental assays. For the sake of simplicity, we assume that division rate and apoptosis rate are constant over time.

#### Estimation of the clone extinction probability

Upon recombination, two daughter cells of different colours are generated and continue to grow, divide and die. After a given time *T* (time of observation), there are four possible outcomes: we can recover the two daughter clones, a single colour clone, green or red (the other clone died before time *T*), and finally both clones disappeared before time *T* (which cannot be measured experimentally). For this estimation, we assumed that both lineages (green and red) have the same proliferation and death rates.

*n*(*t*) is the size (number of cells) of a clone at the time *t*, and it obeys to? a stochastic process that follows the same probability function for every clone. For every clone, we have *n*(0) = 1. The probability of clone disappearance before time *t*. *i.e*. the extinction probability, *p*_0_(*t*) is defined as

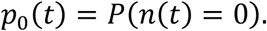

Experimental data gives access to the number of single clones *N_s_* and to the number of twin clones *N_d_* that we observe at the time *T*. From this, we can estimate the probability of extinction at the observation time *T, p*_0_ = p_0_(*T*).

The expected number of single clones is *N_s_* = 2*p*_0_(1 – *p*_0_)

The expected number of twin clones is *N_d_* = (1 – *p*_0_)^2^

which gives the estimator of *p*_0_,

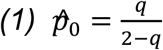

where 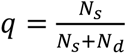

#### Estimation of the clone size and relationship with apoptosis and proliferation rate

Let *μ*(*t*) be mean value of *n*(*t*), clone size in number of cell at time *t*. In theory,

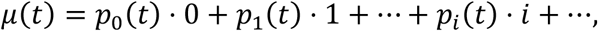

where *p_i_*(*t*) is a probability to observe a clone of size *i* at the given time *t*, or *p_i_*(*t*) = *P*(*n*(*t*) = *i*) and ∑*p_i_*(*t*) = 1 for all *t*. In order to estimate *μ*(*t*) from the observations we use:

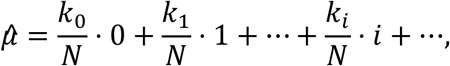

where *N* is a total number of clones (including clones of size “0”), *k_i_* is a number of clones of size *i*, and ∑*k_i_* = *N*.

However, since we cannot observe clones of size 0 we cannot measure *k*_0_ nor *N*. We therefore use 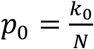 (the probability of clone extinction, see above), and the total number of observed clones *N_obs_* = *N* – *k*_0_. From these two we can estimate 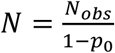. Hence, we estimate *μ* with

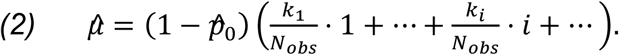

Each clone observed is a result of a stochastic birth-death process, that is fully described by its death rate, *a*, which is a probability of an individual cell to die per unit time and its birth rate, *b*, which is a probability of an individual cell to divide per unit time. While the size of the clone at the given time, *n*(*t*), is a stochastic process, the mean size *μ*(*t*) can be approximated by a deterministic process and is given by the following exponential function:

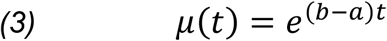

Furthermore, the probability to be extinct at the time *t* (*n*(*t*) = 0) can be expressed as follows [45]:

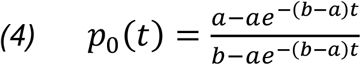

#### Estimating the apoptosis and proliferation rates for each compartment

From the twin clone experimental data, we retrieve for each compartment the proportion of single clone occurrence (**Figure 2C**) and the averaged clone size (**Figure 3A**) which, together with *(1), allows us to* estimate *p*_0_, and *μ* for each of the 40 compartments of the wing disc. These estimates can then be used to estimate parameters *a* and *b* (apoptosis and division rate). For the fixed *t*, based on *(3)* and *(4)*we have:

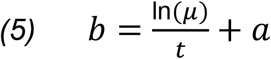

and

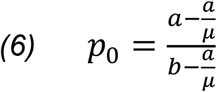

Hence

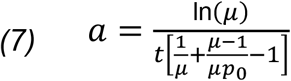

#### Estimating the expected differences of clone size between compartments assuming constant proliferation rate

We use the fixed value of a birth rate, *b* = 0.0657 (average of the estimated *b* from *(5)*), as a fixed value for every compartment. We look for a death rates *α* for each compartment using the expression for *p*_0_(α) (*4*). To the best of our knowledge it is not possible to express explicitly *α* from the given formulation, so we use numerical approach to calculate it. We plot function of *p*_0_ as *p*_0_(*α*) and look for the point of intersection with the measured value 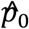. In this way we estimate value of *α*. Then we use *α* and the fixed value of *b* to predict *μ* (*6*), that we then compared with the estimated value 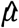 from the experimental data *(2)* correcting for the non-observable disappeared clones (see **Figure 3B**). We overall found a good correlation between the estimated clone size and the prediction, (correlation coefficient *ρ* = 0.6, **Figure 3D**) suggesting that the spatial differences in apoptosis are to a good approximation sufficient to explain the spatial differences in average clone size.

### Statistics

Data were not analysed blindly. No specific method was used to predetermine the number of samples. The definition of n and the number of samples is given in each figure and associated legend. Error bars are standard error of the mean (s.e.m.) or confidence interval 95%.

---

**Supplementary Figure 1 (associated with Figure 1).**
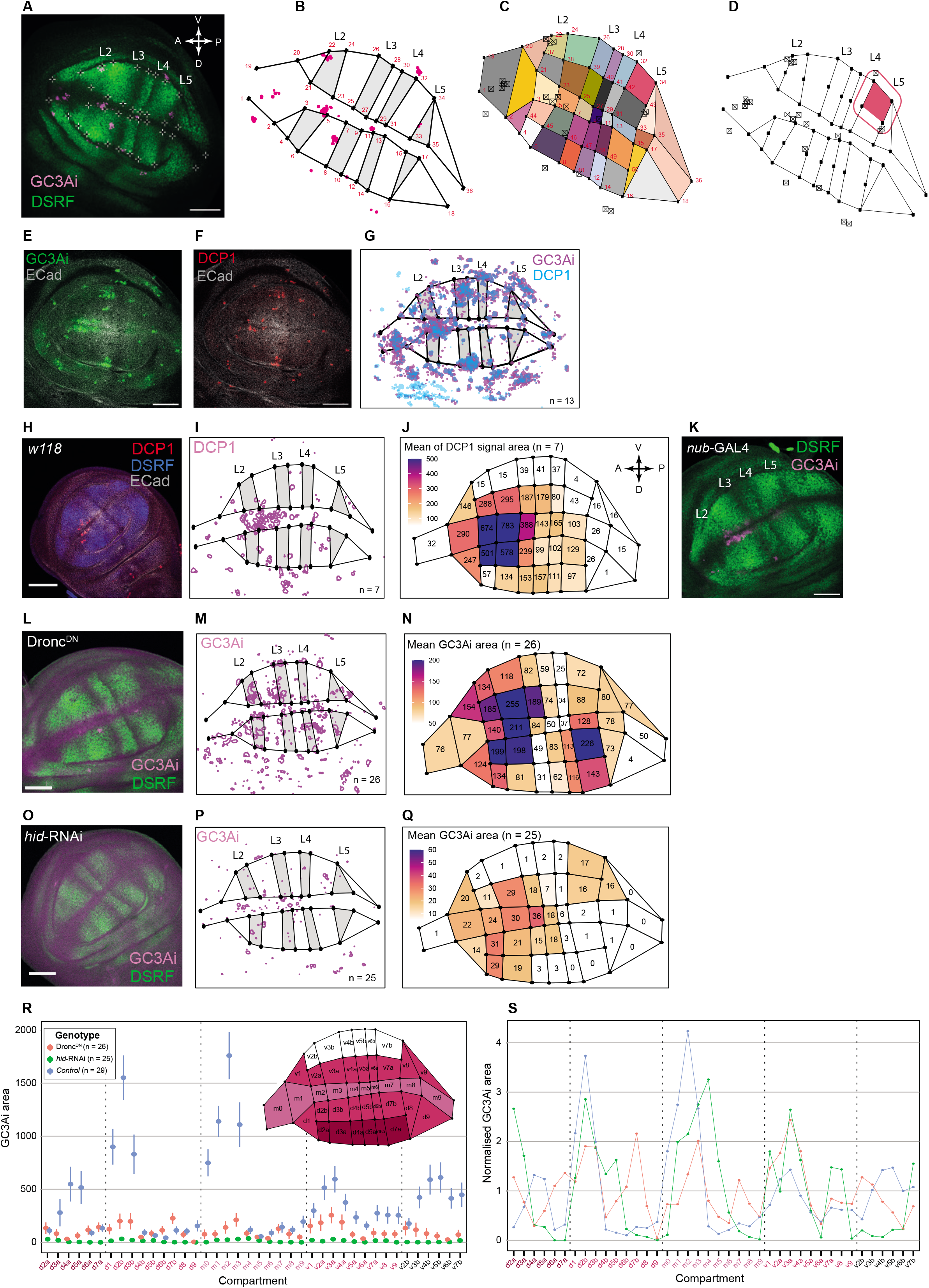
**A:** Wing disc from **Figure1 A** showing the 36 landmarks (white crosses) that were manually positioned at the intersections between veins (L2 to L5), folds and margin of the tissue, visible thanks to DSRF (green). **B:** Map of the wing disc shown in **A**, drawn from the landmark positions (black dots, numbered 1 to 36), and showing the segmented GC3Ai signal (magenta). Vein regions are in grey and intervein/margin regions in white. **C:** Division of the wing disc map shown in **B** into 40 compartments based on polygons (plain colours) defined by adding 14 additional landmarks. (see **Methods**). Boxed black crosses show the centroid of the GC3Ai segmented signal shown in B. **D:** Illustration of the error margin of the compartment boundaries used to account for uncertainty in signal localisation and landmark position. For each compartment, the margin was extended by 11.8 μm to encompass surrounding regions. In this example, the compartment defined by landmarks 32;34;43;42 (coloured pink) has its margin extended and as a result the three GC3Ai centroids located around the compartment (boxed black crosses) will be considered as being also part of this compartment. **(E-G):** Co-localisation of GC3Ai and cleaved DCP-1 (*Drosophila* caspase3). **E** and **F** show the same wing disc with GC3Ai signal (**E**) or anti cleaved DCP-1 immunostaining (**F**). **G:** Superimposition of GC3Ai (magenta) and cleaved DCP-1 (blue) signals from 13 discs. The 13 discs were superimposed using the General Procrustes Alignment (GPA) based on the 36 landmarks positions. **H:**Wings disc from *w118* genotype stained for anti cleaved DCP-1, anti DSRF and anti E-cad. **I:** Superimposition of the segmentation data from 7 discs scaled, rotated and aligned using the GPA approach, showing a pattern for cleaved DCP-1 (magenta) similar than the one for GC3Ai (**Fig. 1F**). **J:** Heat-map showing the average cleaved DCP-1 positive area on each of the 40 compartments of the wing disc in *w118* genetic background. Numbers within each compartment show the average value for cleaved DCP-1 positive area. These values are also shown as a colour coded heat-map. The pattern is similar to the one of GC3Ai (**Figure 1G**). **K:** Wing disc stained with anti DSRF expressing GC3Ai under the control of *nubbin-GAL4*. **(L-N)**: Pattern of GC3Ai in *UAS-Dronc^DN^* background driven by GMR11F02-GAL4, single representative wing disc (**L**), superimposition of 26 discs using GPA (**M**), and with a heat map showing the average GC3Ai signal area over 26 discs (**N**). **(O-Q)**: Pattern of GC3Ai in *hid-RNAi* background driven by GMR11F02-GAL4, single representative wing disc (**O**), superimposition of 25 discs using GPA (**P**), and heat map showing the average GC3Ai signal area over 25 discs (**Q**). **R:** Mean GC3Ai signal area for the 40 compartments in three genotypes (*Control, hid-RNAi, Dronc^DN^*). The inset shows a wing disc diagram with delimitation and names of the 40 compartments on which the quantifications were made. Error bars show s.e.m. **S:** This plot shows the same data than in **R**, but that were normalised to show within each genotype how much the GC3Ai signal varies in a given compartment relatively to the average amount over all the disc. The plot was obtained by dividing the mean value of each compartment (shown in **R**) by the mean over all the compartment means within each genotype.

**Supplementary Figure 2 (associated with Figure 2).**
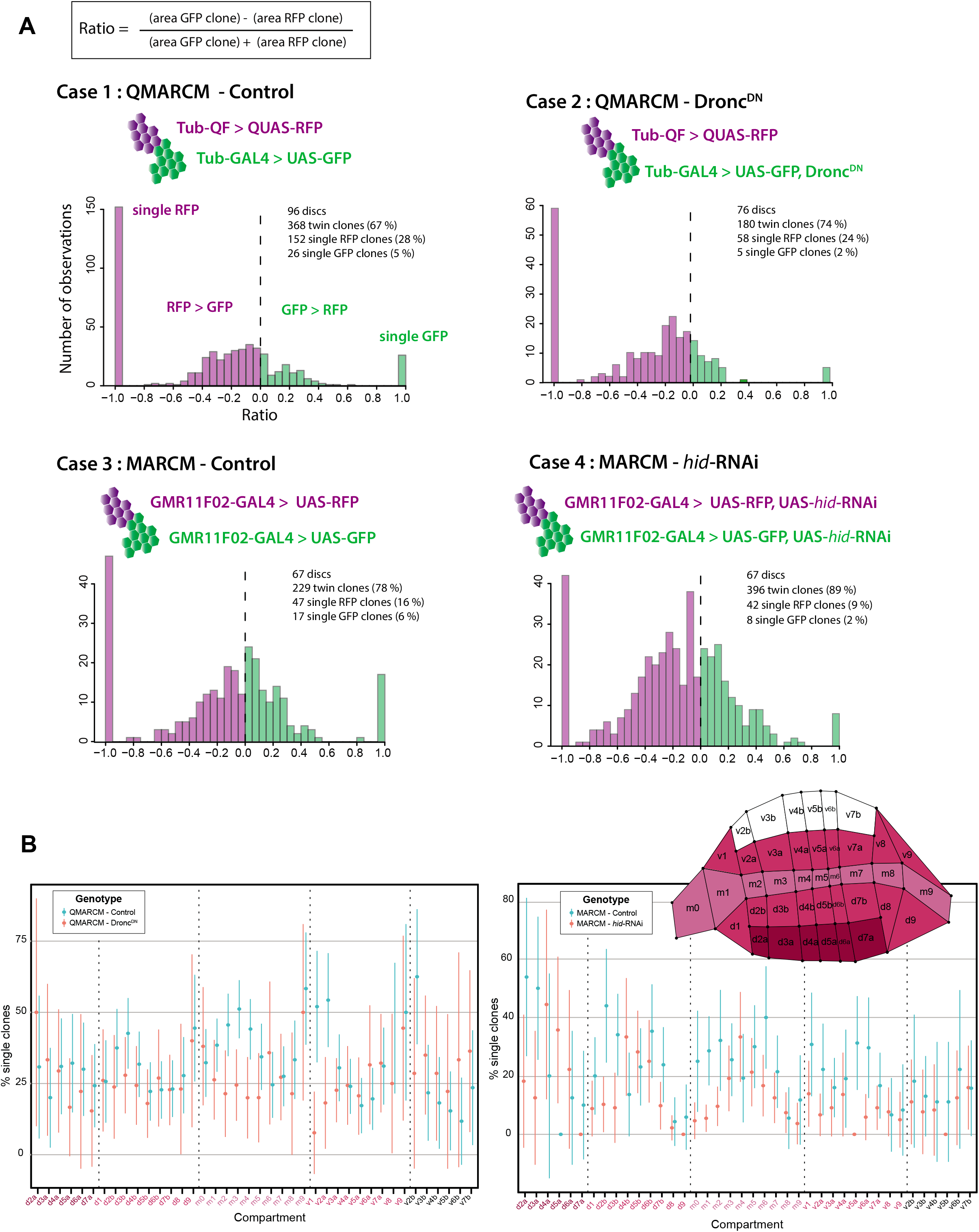
**A:** Distribution of twin-clone Ratio values for each of the four genetic conditions studied to establish the spatial map of single clone occurrences. For each mitotic recombination event, the computed ratio is the difference between areas of the green and red clones over the sum of these areas. Negative ratios indicate twin clones for which the red clone is bigger than the green one whereas twin clones where the green clone is bigger than the red one have positive values. Ratios values of −1 and +1 indicate mitotic recombination events where, respectively, the green or the red clone were lost and have thus a null area. **B:** Frequency of the single colour clones for the 40 compartments in four genotypes (left, QMARCM, QMARCM Dronc^DN^, right, twin spot MARCM, twin spot MARCM *hid*-RNAi). The inset shows a wing disc diagram with delimitation and names of the 40 compartments on which the quantifications were made. Error bars are 95% confidence interval (CI), with 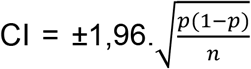, with p, frequency of the single colour clones and n, number of mitotic recombination events observed.

**Supplementary Figure 3 (associated with Figure 3).**
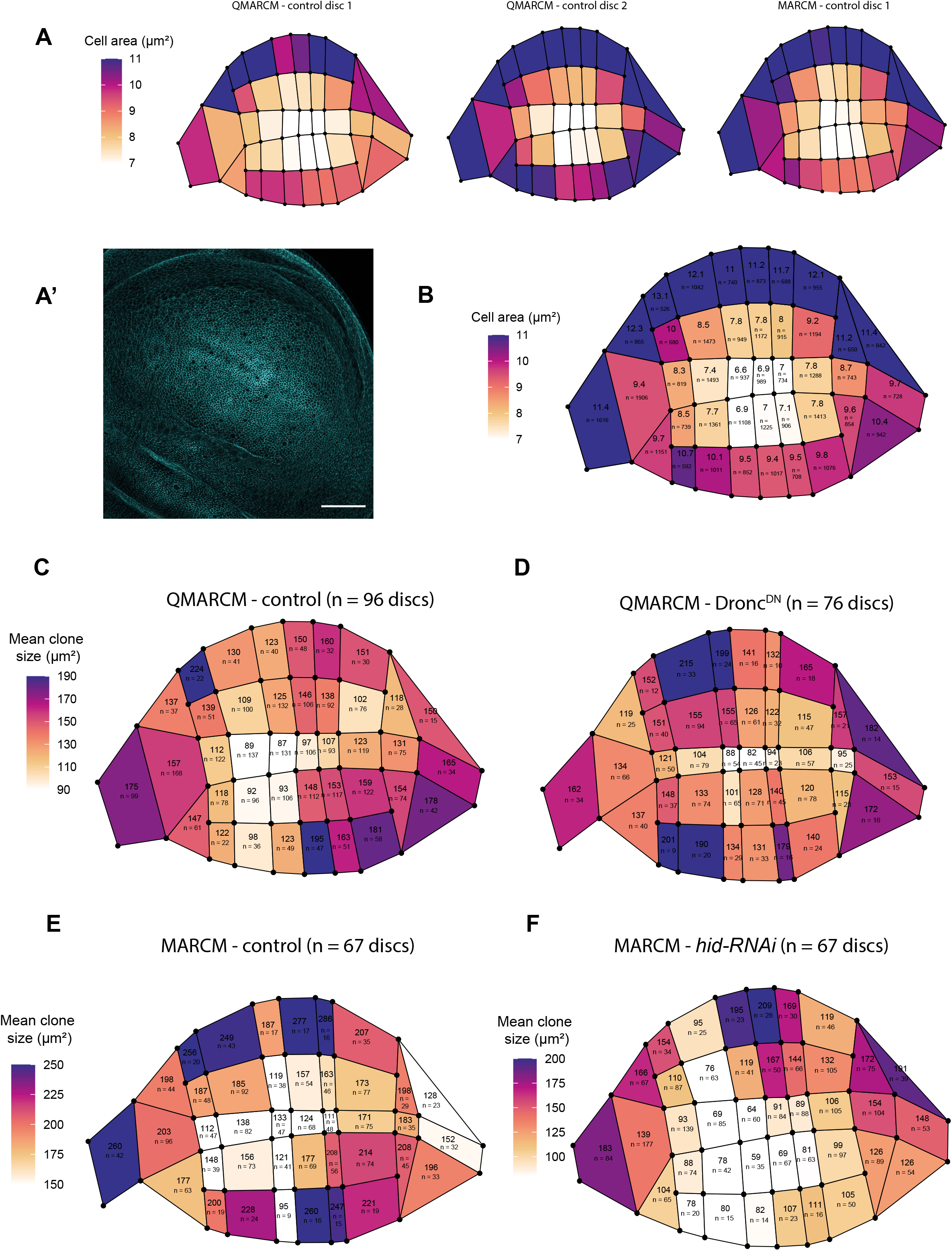
**A:** Heat maps showing the spatial pattern for cell apical area for three segmented discs estimated after local projection of E-cad staining. Two discs were of the *QMARCM – control* genotype and another one from the twin spot *MARCM – control* genotype. The colour code shows the averaged apical cell area for each compartment (same colour scale bar for the three discs).**A’:** Example of a local projection of a wing disc stained with E-cad (the one shown in **A**), allowing the extract cell apical area. Scale bar: 50 μm. **B:** Heat map showing the average spatial pattern of cell size from the three discs shown in A. Within each compartment, the value for mean cell size (μm^2^) and the number of cells measured (over the three discs) are given. **C-F:** Heat maps showing the average spatial pattern for clone surface area (μm2, based on local projections performed at the level of adherens junctions). Each clone was considered irrespectively of its colour and its occurrence as single clone or in a twin spot. For each compartment, the average surface area of the clones and the number of clones examined (n) are given. The data are shown for the four conditions used in this study: QMARCM system, in control conditions (**C**) and upon expression of UAS-Dronc^DN^ in the GFP clone (**D**); twin spot MARCM system driven by the GMR11F02-GAL4, in control condition (**E**) or by expressing UAS-*hid*-RNAi in all the pouch (**F**).

**Supplementary Figure 4 (associated with Figure 4).**
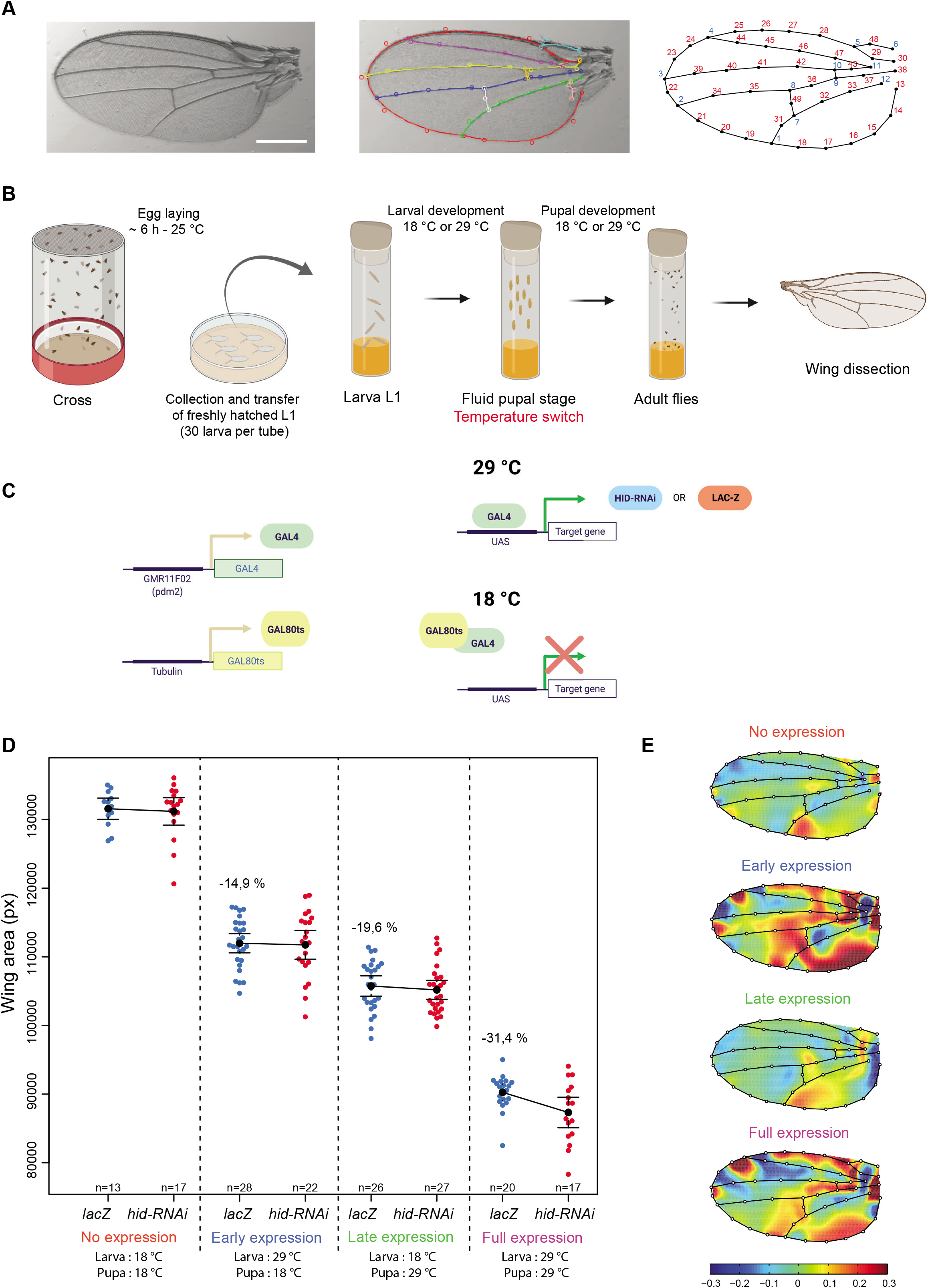
**A:** Adult wing segmentation pipeline. Left: wing example. Middle: the software Wings4 fits 9 spline curves (i.e., equations that give the location of the curve at any point between its end points) to veins and edges of the wing. Each spline is shown with a specific colour. Circles denote the positions of control points, which can be dragged manually by the user to maximise the overlap between the spline curves and veins and edges of the wing. Right: the program CPR extracts from the spline curves the positions of 12 landmarks (blue, numbered 1-12) and 36 semi-landmarks (red, numbered 13-48). Landmarks are strongly reliable points because they are located at the end of the curves. Semi-landmarks are less reliable because they are sampled along the curves by spacing them equally along each curve segment. **B:** Experimental design for conditional gene expression during early or late developmental stages. Flies were crossed in cages supplied with agar petri dishes and allowed to lay eggs for ~ 6 h at 25 °C. Freshly hatched larva were transferred to regular vials at a density of 30 individuals / tube, and placed at 18 °C or 29 °C. Once arrived at the fluid pupal stage, individuals were switched temperature (from 18 °C to 29 °C and vice-versa). Upon hatching flies were collected for wing dissection. **C:** Rationale for conditional gene expression using tub-GAL80^ts^. GAL80^ts^ has ubiquitous expression under the control of tubulin promoter whereas GMR11F02-GAL4 expression is restricted to wing tissue. At 18 °C, GAL80^ts^ binds to the GAL4 and inhibits its activator function, precluding the expression of the target genes. At 29 °C Gal80^ts^ is unable to bind GAL4, allowing GAL4 binding to the upstream activating sequence (UAS) and transcription of target genes (*lacZ* or *hid-RNAi*). **D:** Variation of adult wing size caused by expression of *hid*-RNAi under the control GMR11F02-GAL4 driver at different developmental stages. The effect of expressing *hid*-RNAi compared to *lacZ* is shown, for four conditions (No expression; Expression before fluid pupal stage (Early expression); Expression after fluid pupal stage (Late expression); Expression during all development (Full expression). Each blue and red dot represents an individual wing size value, black dots are means, and error bars are 95 % confidence interval of the mean. For each condition, the percentage of variation of the *lacZ* mean relative to the “No expression” condition is shown. Pairwise t-test for *lacZ* compared to *hid-RNAi* within each condition are all non-significant. **E:** Local variations of shape caused by expressing *hid*-RNAi at different developmental stages. Colours represent changes in relative area necessary to transform the average wing for *lacZ* group into the average wing of *hid*-RNAi group (red: expansion, blue: contraction). Colour bar shows the upper and lower limits in deformation. Highest colour intensity is reached at 30 % increase and decrease in local area in *hid*-RNAi group with respect to *lacZ* group. Panels B and C were created with BioRender.com.

